# Sero-prevalence of anti-SARS-CoV-2 Antibodies in Addis Ababa, Ethiopia

**DOI:** 10.1101/2020.10.13.337287

**Authors:** Berhanu Nega Alemu, Adamu Addissie, Gemechis Mamo, Negussie Deyessa, Tamrat Abebe, Abdulnasir Abagero, Wondimu Ayele, Workeabeba Abebe, Tewodros Haile, Rahel Argaw, Wondwossen Amogne, Ayele Belachew, Zelalem Desalegn, Brhanu Teka, Eva Kantelhardt, Mesfin Wossen, Saro Abdella, Getachew Tollera, Lia Tadesse

**Affiliations:** Department of Surgery, School of Medicine, College of Health Sciences, Addis Ababa University, Addis Ababa, Ethiopia; Department of Preventive Medicine, School of Public Health, College of Health Sciences, Addis Ababa University, Addis Ababa, Ethiopia; Mulu-G-Health Services, Addis Ababa, Ethiopia; Department of, Microbiology, Immunology and Parasitology, School of Medicine, College of Health Sciences, Addis Ababa University, Addis Ababa, Ethiopia; Department of Pediatrics and Child Health, School of Medicine, College of Health Sciences, Addis Ababa University, Addis Ababa, Ethiopia; Department of Internal Medicine, School of Medicine, College of Health Sciences, Addis Ababa University, Addis Ababa, Ethiopia; Institute for Medical Epidemiology Biometrics and Informatics, Martin Luther University, Halle-Wittenberg, Germany; Ethiopian Public Health Institute, Ministry of Health of Ethiopia, Addis Ababa, Ethiopia; Ministry of Health of Ethiopia, Addis Ababa, Ethiopia

**Keywords:** sero-prevalence, Ethiopia, Addis Ababa, SARS-CoV-2, Antibody Testing

## Abstract

**Background:** Anti-SARS-CoV-2 antibody tests are being increasingly used for sero-epidemiological purposes to provide better understanding of the extent of the infection in the community, and monitoring the progression of the COVID-19 epidemic. We conducted sero-prevalence study to estimate prior infection with with SARS-CoV-2 in Addis Ababa.

**Methods:** A cross-sectional study was done from April 23 to 28, 2020 among 301 randomly selected residents of Addis Ababa; with no known history of contact with confirmed COVID-19 person. Interviews on socio demographic and behavioural risk factor followed by serological tests were performed for SARS-CoV-2 IgM, and IgG antibodies, using COVID-19 IgG/IgM Rapid Test Cassette. The test has sensitivity of 87·9% and specificity of 100% for lgM; and a sensitivity of 97·2% and specificity of 100% for IgG. RT-PCR test was also done on combined nasopharyngeal and oropharengeal swabs as an important public health consideration.

**Findings:** The unadjusted antibody-based crude SARS-CoV-2 prevalence was 7·6% and the adjusted true SARS-CoV-2 prevalence was estimated at 8·8% (95% CI 5·5%-11·6%) for the study population. Higher sero-prevalence were observed for males (9.0%), age below 50 years (8.2%), students and unemployed (15.6%), those with primary education (12.1%), smokers (7.8%), alcohol consumers (8.6%), chatt-chewers (13.6%) and shish smokers (18.8%). Seroprevalence was not significantly associated neither with socio-demographic not behavioral characteristics. According to the findings, possibly more individuals had been infected in Addis Ababa than what was being detected and reported by RT-PCR test suggestive of community transmission. The use of serological test for epidemiological estimation of the extent of SARS-CoV-2 epidemic gives a more precise estimate of magnitude which would be used for further monitoring and surveillance of the magnitude of the SARS CoV-2 infection.

## Introduction

Testing SARS-CoV-2 is made in two ways, by detecting the virus itself (RT-PCR) and by detecting the host’s response to the virus (serology)^1^. The World Health Organisation (WHO) recommends RT-PCR test as gold standard for COVID-19 case identification^2–4^. The clinical course of SARS-CoV-2 infection ranges from asymptomatic to fatal^5, 6^ making the identification of “cases” very complex. The response plan to the pandemic requires epidemiological data that shows the true magnitude of the disease and the serology test can be used for such purposes^7^. The specificity of the RT-PCR test is almost 100%. However, sensitivity is reported as low as 64%^8–10^. Differences in the detection limits of the test kits used, low initial viral load due to timing as well as improper specimen collection^11–13^ may account for such low sensitivity.

The lower sensitivity or “false negative” RT-PCR results and/or the “immune passport” led to the rapid development of serologic assays^14–16^. The serologic tests in the market have different formats (lateral flow immunoassays, ELISA, and chemiluminescent immunoassays), detect different classes of antibodies (total antibody, IgG, IgM, and IgA), target different antigens (recombinant nucleocapsid protein [NP], subunit 1 of the spike glycoprotein [S1], and the Spike glycoprotein receptor-binding domain [RBD]) and accept different types of the specimen (whole blood, serum, and plasma)^16^. Data generated from population-based serology can be used for various purposes including estimation of community transmission rates and assess the impact of non-pharmacological interventions^14^. Despite the potential of the serology test, equally, the availability of a serology test with excellent sensitivity and specificity is a discussion point.

Since the report of the first confirmed SARS-CoV-2 inflection in Ethiopia on March 13, 2020, the rate of increment has been largely slow at the beginning. This required a more precise estimation of the magnitude of SARS CoV-2 infection on top of the estimates based on routine RT-PCR testing. To this end, we set a goal to conduct a series of sero-epidemiological studies. Here were reporting our first findings on anti-SARS-CoV-2 antibody among permanent residents in Addis Ababa conducted to estimate what extent SARS-CoV-2 infection has been circulating in the population.

## Methods

### Study design and participants

A cross-sectional community-based study was conducted from April 23-28, 2020. The study was undertaken in Addis Ababa, the capital city of Ethiopia, with an estimated total population of 4,793,699^17^.

The study population were adult (≥ 18 years), residents of Addis Ababa, with no history of known contact with confirmed COVID-19 cases and no recent history of travel out of Ethiopia in the past four months prior to the period of data collection. Individuals who were unable to consent, those with unstable mental state and suspected to have acute SARS-CoV-2 infection were excluded.

Sample size was calculated using a samples size formula for single population proportion, assuming the proportion of people with immunity among community members. with no close contact with SARS-CoV-2 Infected individuals to be14% based on earlier European studies about the time of the study ^18^, 80% power and 99% confidence level The Epi Info/Open Epi program is used for the sample size calculation^19^. Accordingly, the sample size calculated was 319.

Study participants were recruited from the community level randomly through sub-city health extension focal personnel. All randomly selected individuals who fulfilled study inclusion criteria were invited to the study. They were approached and received an invitation through health extension workers (HEWs) from the respective sub-city. All study participants were briefed about the study and gave written consent before proceeding to interview and sample collection.

### Data collection

Socio-demographic and clinical data were collected through interviewer administered questionnaire. Whole blood sample for antibody testing and combined nasopharyngeal and oropharengeal swab for RT-PCR testing was collected from each participant. A questionnaire comprising socio-demographic characteristics such as age, sex, marital status, and place of residence; a history of illness in the past three to five months like signs and symptoms of pneumonia and/ or influenza-like illness; and perceived severity of illness, etc was administered. Prior to data collection, the questionnaire was pre-tested and necessary adjustments made. Peripheral blood (4ml) was collected from each participant by experienced phlebotomist by venipuncture using Ethylenediaminetetraacetic acid (EDTA) tubes and transported to the Ethiopian Public Health Institute (EPHI) Influenza laboratory for analysis. Plasma was extracted following centrifugation at 3000rpm for 10 minutes in level-II bio-safety cabinet. Plasma was stored using cryovials in refrigerator at 2-8°C until the next day. The leftover plasma was stored at −80°C for future use at EPHI bio-bank. A combined nasopharyngeal and oropharengeal swab was collected from each participant using a Dacron or polyester flocked swabs (*KANGJIAN Medical* Apparatus *Co*., *Ltd*.) by trained personnel, and in Viral Transport Medium (VTM) (Longsee, LAKEbio) transported to EPHI for RT-PCR analysis of SARS-CoV-2 in cold chain (2-8°C) and processed immediately or stored for 1 to 3 days at −80°C.

### Laboratory Analyses of collected samples

#### Serology

Analysis for the presence of plasma IgM and/or IgG antibodies was done using a commercially available immunolateral flow immunoassay kit (Zhejiang Orient Gene Biotech Co Ltd, Huzhou, Zhejiang, China) following manufacturers instruction. The interpretation of the test was made by two experts (microbiologists and laboratory technologist). According to the manufacturer the test has a sensitivity and specificity of 87.9 and 100% respectively. Another validation study has reported the test has lower sensitivity (69% for IgM and 93.1% for IgG and the specificity was found to be 100% for IgM and 99.2% for IgG using RT-PCR assay as a comparator^20^.

#### RNA extraction and RT-PCR analysis of COVID-19

The combined swab (Nasopharyngeal and oropharyngeal) was transferred into lysis buffer that contains a guanidinium-based inactivating agent and viral RNA was extracted using Nucleic Acid Isolation Kit (Da’an Gene Corporation) following manufacturer’s instruction. Briefly a 200 μL of combined swab in VTM was used for viral RNA extraction and viral RNA was eluted with 60 μL elution buffer. Real-time reverse transcriptional polymerase chain reaction (RT-PCR) reagent of Da’an Gene cooperation was employed for SARS-CoV-2 detection following manufacturer’s protocol. Briefly, two PCR primer and probe sets, which target the open reading frame 1a/b (ORF 1a/b) (FAM reporter) and nucleocapsid protein (N) genes and N (VIC reporter) genes were added in the same reaction mixture. In each run, positive and negative controls were included. Samples were considered to be positive when both sets gave reliable signal (<40CT value).

#### Data processing and Analysis

All demographic, epidemiologic and laboratory data were cleaned and entered into EPI DATA using a controlled and programmed data entry format. The data was coded and anonymized data were merged to laboratory data which were exported into SPSS for windows. Descriptive statistics was used to summarise key findings. Estimation of true values for sero-prevalence for the given levels of sensitivity and specificity were computed via web-based statistical software and epidemiologic calculator including EPI Tools (https://epitools.ausvet.com.au/trueprevalence)^21^. Binary logistic regression was used to identify factors associated with the outcome variable. Variables whose p value less than 0.05 at bivariate analysis were included in multivariable analysis.

#### Ethical Considerations

The research proposal is approved by the Institutional Review Boards (IRB) of both the College of Health Sciences of the Addis Ababa University and EPHI. Written informed consent was obtained from all study participants. All study participants were informed of their RT-PCR test results as per the national testing protocol.

#### Funding

While we had no any official sponsor, costs of the research were met by resources from Addis Ababa University and the Ethiopian Public Health Institute (EPHI).

## Results

### Characteristics of the study participants

A total of 301 participants were included in the study with a response rate of 94.4%. Most of the study participants were in the age group 18-30 years (48.2%) (with a median age of 30 years ± 10.9 years), males (62·5%), single (47·0%), and health professionals (25·6%), educated above high school (37·9%), non-smokers (78·7%), with no history of regular alcohol (42·5%), chat (70·8%), shisha (94·7%) use, vaccinated for BCG (47·8%), No contact with PCR confirmed COVID-19 cases (99·0%) (Table1).

**Table 1:** Socio demographic and behavioural characteristics of participants in Addis Ababa Ethiopia, May, 2020 (n=301)

| Socio-demographic and behaviour factors |  | Number | Percent |
| --- | --- | --- | --- |
| Sex | Male | 188 | 62.5% |
|  | Female | 113 | 37.5% |
| Age group | 18-29 | 145 | 48.2% |
|  | 30-49 | 122 | 40.5% |
|  | above 49 | 34 | 11.3% |
| | Median age = 30 years ± 10.9 years | | |
| Educational status | No formal education | 16 | 5.3% |
|  | 1-4 grade | 16 | 5.3% |
|  | 5-8 grade | 75 | 24.9% |
|  | 9-12 grade | 80 | 26.6% |
|  | 12 and above | 114 | 37.9% |
| Current marital status | Married | 133 | 42.9% |
|  | Single | 132 | 47.0% |
|  | Divorced//widow | 36 | 12.0% |
| Occupation | Health care worker | 77 | 25.6% |
|  | Driver/parking/mechanics | 8 | 2.6% |
|  | Bank/ supermarket/ shop/Hotel | 8 | 2.6% |
|  | Daily labourers/ Petty trading | 114 | 37.9% |
|  | (Government/NGO) | 48 | 15.9% |
|  | (Housewife /Retired) | 15 | 5.0% |
|  | Student/ Unemployed | 31 | 10.3% |
| Smoked a cigarette | No | 237 | 78.7% |
|  | yes | 64 | 21.3% |
| Consumed alcohol | No | 162 | 53.8% |
|  | yes | 139 | 46.2% |
| Chewed chat | No | 213 | 70.8% |
|  | yes | 88 | 29.2% |
| Used shisha | No | 295 | 94.7% |
|  | yes | 6 | 3.3% |

### SARS-CoV-2 Sero-prevalence

Out of the 301 individuals included in the study, 23 (7·6%) tested positive for anti-SARS-CoV-2 antibodies with an unadjusted antibody-based crude prevalence was 7·6%. Accordingly true prevalence adjusted for the test sensitivity and specificity was estimated at 8·8% (95% CI 5·5%-11·6%). Higher sero-prevalence were observed for Males (9.0%), age below 50 years (8.2), students and unemployed (15.6), those with primary education (12.1), smokers (7.8), alcohol consumers (8.6), chatt-chewers (13.6%) and shish smokers (18.8%). (Table 2)

**Table 2:**
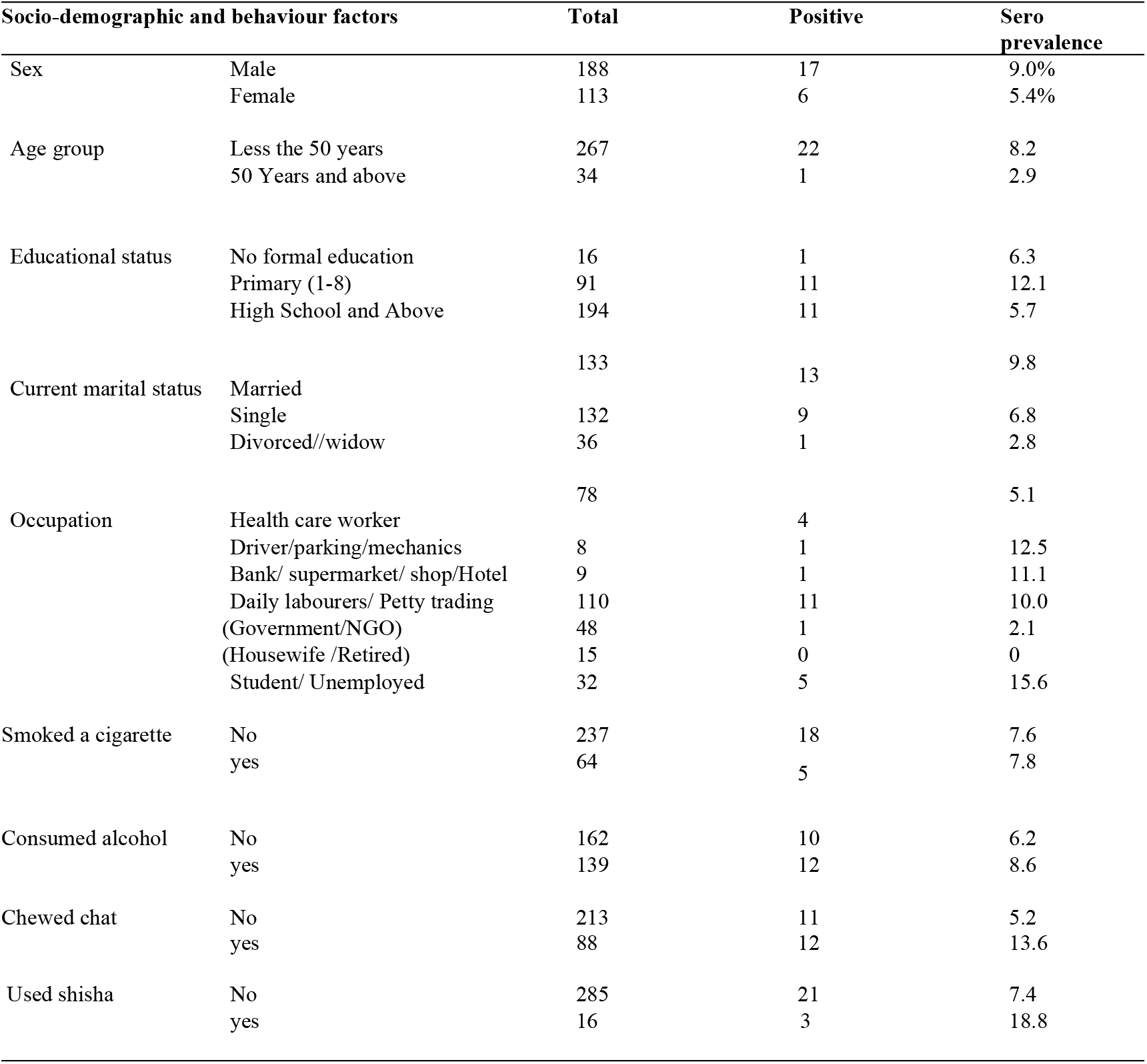
Sero-prevalence study participants by demographic and behavioural characteristics, Addis Ababa Ethiopia, May, 2020.

### Factors Associated with SARS-CoV-2 Seropositive tests

On binary logistic regression there was negative association between chat chewing and SARS Co2 Seropositivity (OR=0·35, 95% CI: 0·2, 0·9). Associations were lost on adjusted logistic regression. . Government/NGO workers were found to be associated as protective effect for SARS-CoV-2 sero status (OR=0·052, 95% CI: 0.0, 0·57) (Table 3)

**Table 3:** Factors Demographic and behavioural characteristics associated with Sero prevalence in bivariate and multivariable analysis in Addis Ababa Ethiopia, May, 2020

| Socio-demographic and behaviour factors |  | Negative |  | Positive |  |  |  |
| --- | --- | --- | --- | --- | --- | --- | --- |
|  |  | Number | Percent | Number | Percent | COR(95CI%) | AOR(95CI%) |
| Sex | Male | 171 | 61.5% | 17 | 73.9% | 0.27(0.03,2.45) | 1.01(0.31,3.26) |
|  | Female | 107 | 38.5% | 6 | 26.1% |  |  |
| Age group | Less the 50 years | 245 | 88.1% | 22 | 95.7% | 2.96(0.39,22.71) | 0.18(0.02,2.07) |
|  | 50 Years and above | 33 | 11.9% | 1 | 4.3% | 1* |  |
| | Median age = 30 years ± 10.9 years | | | | | | |
| Educational status | No formal education | 15 | 5.4% | 1 | 4.3% | 1.11(0.13,9.18) | 3.47(0.28,42.89) |
|  | Primary (1-8) | 80 | 28.7% | 11 | 47.8% | 2.29(0.95,5.50) | 1.73(0.54,5.58) |
|  | High School and Above | 183 | 65.8% | 11 | 47.8% | 1* |  |
| Current marital status | Married | 120 | 43.2% | 13 | 56.5% | 3.79(0.48, 30.00) | 4.56(0.47,44.73) |
|  | Single | 123 | 44.2% | 9 | 39.1% | 2.56(0.31, 20.91) | 2.09(0.21,21.32) |
|  | Divorced//widow | 35 | 12.6% | 1 | 4.3% | 1* |  |
| Occupation | Health care worker | 74 | 26.6% | 4 | 17.4% | 0.29(0.073,1.17) | 0.29(0.06, 1.34) |
|  | Driver/parking/mechanics | 7 | 2.5% | 1 | 4.3% | 0.77(0.077,7.71) | 0.15(0.01, 1.99) |
|  | Bank/ supermarket/ shop/Hotel | 8 | 2.9% | 1 | 4.3% | 0.66(0.069,6.65) | 0.36(0.03, 3.99) |
|  | Daily labourers/ Petty trading | 99 | 35.6% | 11 | 47.8% | 0.60(0.192,1.88) | 0.22(0.05, 1.02) |
|  | Government and NGO | 47 | 16.9% | 1 | 4.3% | 0.11(0.013,1.03) | **0.052(0.01, 0.57)** |
|  | Student/ Unemployed | 27 | 9.7% | 5 | 21.7% | 1* |  |
| Smoked a cigarette | No | 219 | 78.8% | 18 | 78.3% | 0.97(0.34,2.72) |  |
|  | yes | 59 | 21.2% | 5 | 21.7% | 1* |  |
| Consumed alcohol | No | 152 | 54.7% | 10 | 45.5% | 0.69(0.29,1.65) | 0.93(0.34,2.55) |
|  | yes | 126 | 45.3% | 12 | 54.5% | 1* |  |
| Chewed chat | No | 202 | 72.7% | 11 | 47.8% | **0.35(0.146,0.815)** | 0.39(0.13,1.18) |
|  | yes | 76 | 27.3% | 12 | 52.2% | 1* |  |
| Used shisha | No | 264 | 95.3% | 21 | 91.3% | 0.52(0.11,2.45) |  |
|  | yes | 13 | 4.7% | 2 | 8.7% |  |  |

## Discussion

As far as our literature review during the writing of this paper goes, this was the first SARS-CoV-2 sero-prevalence study in Ethiopia which sheds light on the extent of SARS-CoV-2 in a time point when there was a range of dilemma reflected on the true extent of the infection in Africa due to the slow trajectory of cases. The findings of the study indicated that there is a proportionally high level of SARS-CoV-2infection in the community compared to the number of reported cases via active surveillance as well as the number of cases projected for the given period. This requires evidence-based explanation.

Viral antigen test (RT-PCR) is primarily done to identify new cases while antibody tests are used for estimation of past infections who already have recovered without possibly realizing it. This helps us to understand the spread of the disease within the community and possibly predict its progression during the coming months. It also helps to identify those who have already been infected and developed immunity^22^. The duration of the exposure depends on the type of antibody. However, it is not known how long the antibodies it last. The duration of the exposure depends on the type of antibody. However, it is not known how long the antibodies it last. Reports have documented that antibody detection offers vital clinical information during the course of SARS-CoV-2 infections with potential application in the diagnosis and management of COVID-19 patients^23^.

While RT-PCR is a gold standard for the diagnosis of SARS-CoV-2 infection, sero-prevalence antibody studies are used to approximate for the local epidemiology of the infection and to monitor disease progression in the community. WHO discourages use of antibody testing for clinical and individual diagnostic purposes, while its use for epidemiologic studies and surveillance purposes is highly encouraged with local validation of available kits^7^.

### Sero-prevalence

The study identified an overall SARS-CoV-2 sero-prevalence of 7·6% which is found to be higher than the estimates based on RT-PCT testing in Addis Ababa, during the period of the study. It is recommended that adjustments are made on the crude estimates of prevalence values which are based on screening tests. To this effect, the true estimate adjusts the values to the levels of sensitivity and specificity of the screening tests employed. The COVID-19 IgG/IgM Rapid Test Cassette screening test used in the study has 87% sensitivity and100% specificity as is reported by the manufacturer. The true value estimate using EPI Tool showed an overall sero-prevalence of 8·8% (95% CI 5·5%-11·6%) Validation conducted for the same test in Sweden has documented relatively lower values of sensitivity (69% for IgM and 93·1% for IgG), while kept the specificity high (100% for IgM and 99·2% for IgG) ^20^. This reflects the fact that the actual estimates of sero-positivity in the study subjects could even be higher, when adjusted to the lower values of sensitivity.

Sero-prevalence studies conducted in other countries during the period of the study reported varied magnitudes of SARS-CoV-2 sero-prevalence: Spain health workers (9·3%)^24^ and community (5%)^18^; Germany (14%)^25^, Switzerland (5·5%)^25^, Massachusetts (30%)^25^, Los Angeles (4·1%)^25^, Santa Clara (1·5%)^25^, New York (13%)^25^, Iran (22%)^26^, and Brazil (4%)^27^. The differences between our findings and the mentioned reports could be explained by technical and methodological provisions of the tests, the risk levels of the tested population and the extent of SARS-CoV-2 infection on the community.

### The magnitude of SARS-CoV-2

The findings signify the importance of documenting the true extent of SARS-CoV-2 infection in Addis Ababa. Based on the official reports of cases in Ethiopia, there were 126 confirmed cases (as of April 28, 2020)^28^ of which the majority are from Addis Ababa. On the other hand, based on SARS-CoV-2 modelling for Addis Ababa, the total number of infected and recovered individuals would have been 430 cases in the same week corresponding to our study. According to the estimates of our study and adjusted true values of 8·8% (95% CI 5·5%-11·6%) the estimated total number of individuals who are possibly infected and recovered would be 421,845 (95% CI, 254,066 - 584,831)^17^. This is way higher than the official RT-PCR report and the projected estimated and released by EPHI^29^.

The possible reasons for the very high levels of estimated numbers of infected and recovered individuals beyond the available estimates could fall into issues around the adequacy of RT-PCR testing, predominantly asymptomatic individuals, and last but not least presence of cases already in the city log before the official reports.

a. Timing of the first cases in Addis Ababa: Similar discrepancies have been documented elsewhere between the number of cases reported and the estimates based on serological testing. Those studies have accordingly concluded that SARS-CoV-2 infection “was there before we knew it”. Even though the first case of SARS-CoV-2was reported on March 13, 2020, as per the findings in the study it is possible that the infection was definitely in the community before the time we knew it. There are anecdotal reports that there have been cases with severe respiratory signs and symptoms in late December to early January about the time when official reports of cases were made in China. Owing to the frequent air traffic between various Chinese cities in Addis Ababa over 10 times per week through Ethiopian airlines makes it difficult to rule out the possibility of such early cases in Addis Ababa. Besides, a review of health facility reports of respiratory illnesses in Addis Ababa showed a two-fold increase in the 6 months period (September 2019 to February 2020), compared to the level in the previous year^30^.
b. The RT-PCR testing issues: If the magnitude reported based on the current serology estimate is closer to the true estimate, then why was it not possible to detect this by active surveillance. It is known that the RT-PCR test detects active cases and not past infections and recovered patients. If individuals were already infected and recovered they will not be detected by the RT-PCR. It is also possible that the RT-PCR misses some active cases due to some factors giving rise to less sensitivity. Apart from the inherent nature of the test, the yield depends on the total number of individuals tested and what population groups are being targeted for testing. In the Ethiopian setting until recently about 70-75% of the cases nationally reported are those returning from other countries. This is followed by the contacts of those individuals. The national testing strategy used to focuse on testing these particular groups and suspected individuals making it almost impossible to detect any possible asymptomatic cases in the community which would require extensive testing of representative targets in the community. On top of that in a community where the majority have been already infected and recovered, we expect less RT-PCR positivity if they already have some level of acquired protection. Subsequent to the communication of the preliminary findings at Ministry of Health, the testing focused more on the community levels and the number of confirmed cases has increased since then.
c. Predominantly asymptomatic cases: Another reason that explains the gap with the sero-prevalence study is to do with the majority of the SARS-CoV-2infection being asymptomatic. According to the widely reported natural history of SARS-CoV-2 infection, the majority (80-85%) will have a mild form of illness. Even though there is no close to exact estimates, there are increasing reports that there are significant portions of asymptomatic or pre-symptomatic SARS-CoV-2 infections. This could be determined by some factors such as age, co-morbidities, immune status and possibly ecological factors to do with the viral strain and virulence. The Addis Ababa population is predominantly young and the prevalence of chronic diseases is relatively low compared to western countries.

In the presence of such a big burden of cases in a big city like Addis Ababa, why did not we see an increase or surge in the number of severe cases in the health facilities? The same factors i.e. predominantly asymptomatic cases could explain this owing to the predominantly young population and lower levels of non-communicable diseases in the Ethiopian population. However, our study did not get any supporting evidence to the looming theory of the protective association between BCG Vaccination and SARS-CoV-2 infection.

### The implications of the study

As the first antibody testing among the general public in Addis Ababa, the study signified the need for extensive serological evidence to make comprehensive estimates of the true extent of SARS-CoV-2 infection in the population especially in a situation where we have the limited testing capacity for RT-PCR. Comparative advantages of antibody testing include cost-effectiveness, easy technology, and point of care testing. This makes serological estimates very much appealing to the low-resource settings. However, in the given circumstances, serological tests are taken as complementary rather than independent in generating epidemiologic evidence for SARS-CoV-2 infection. This is indicative of the need for more precise population-based sero prevalence and surveillance to have a better sense of the extent of the epidemic and to monitor its progress over time, which further guides the design and implementation of targeted interventions. As this is the first of series of studies we planned, we plan to conduct nationwide sero-prevalence study and possibly set-up sentinel sero-prevalence longitudinal studies.

### Policy implications

Such serological estimates provide a substantial and comprehensive input to the epidemiologic projections of SARS-CoV-2 especially in a setting where extensive and reliable viral antigen tests are a limitation. In addition, the current and subsequent data will provide evidence for policy makers to make decisions on locally generated evidence.

While this possibly the first SARS-CoV-2 sero-prevalence study in Ethiopia (possibly in Africa, there are a number of limitations. The test kit utilized was not validated in our population and the target antigen of the kit is not described by the company. In addition the relatively smaller number subjects included in the study may affect generalisability. The test kit has been validated in Sweden and used for population based studies in Spain and China. In-country validation of the study is currently being done by EPHI but not completed. The serological tests were not primarily done to detect a concurrent active infection with no contribution to SARS-CoV-2 case detection and management purposes.

## Conclusion

- The extent of sero-prevalence indicates that more individuals are infected by SARS-CoV-2 in Addis Ababa more than what is being detected and reported by RT-PCR testing, as well as what was estimated by current prediction models.
- Based on analysis of projections and estimates, it shows that there could have been SARS-CoV-2 infections in Ethiopia earlier before the first case reported in March 13^th^
- Majority of SARS-CoV-2infections in Addis Ababa are primarily either mild or asymptomatic contributing to the less identification of cases
- The sero-prevalence results indicate community transmission has already been established in Addis Ababa

## Recommendations

- The approach would serve as a base-line and a model of analysis for further serological investigation and its application in Ethiopia and other African countries.
- It is recommended that a more extensive sero-prevalence study be conducted with a more representative population sample to further establish more substantial evidence on the extent of SARS-CoV-2infection in Addis Ababa and nationally.
- Establish a system for longitudinal sero-prevalence studies in Addis Ababa and other parts of the country to have a continued precise estimate of the extent of SARS-CoV-2infection
- The national estimate of cases should be complemented with appropriate serology tests in the community to diagnose and identify levels of community transmission and guide containment strategies among different population groups while maintaining non-pharmacological interventions

## Contributors

BN, GM, AB, AdA and ND contributed to the conceptualization of the study. TA, AbA, WAy, TH, RA, ZD, EK, MW, SA, GT, and LT contributed to further developments of the research idea and study design. BN, AdA, ND, TA, AbA, WAb, GM, WAm, Way, TH, RA, AB, ZD, BT, GT, MW, and LT were involved in the organization of data collection, data analysis and interpretations of the findings. AdA, BN, ND, AbA, TA, ZD, WAy, WAm and WAb were involved in drafting the initial version of the manuscript. All authors were involved in writing the final version of the manuscript.

## Data sharing

Addis Ababa University, Ethiopian Public Health Institute, and the Federal Ministry of Health dataset can be accessed by researchers. Ethical approvals from the IRB of the College of Health Sciences and the IRB of the Ethiopian Public Health Institute are available. Researchers wishing to directly analyze patient-level data will be required to request the corresponding author. Patient-level data cannot be taken out of the secure network.

## Acknowledgments

We would like to acknowledge the Office of the Prime Minister of Ethiopia, Office of Ministry for Health of Ethiopia, Addis Ababa Regional Health Bureau, and Addis Ababa University for the various levels support they provided. Dr Kjell Magne Kiplesung from Nordic Medical Centre provided the test kits and the Ethiopian Public Health Institute allowed the laboratory for tests.

